# A region-specific murine intestinal monolayer platform for assessing iron form–dependent transepithelial transport

**DOI:** 10.64898/2026.05.09.717085

**Authors:** Yuta Takase, Yoshiko Murata, Kosuke Namba, Toshio Takahashi

**Affiliations:** Suntory Foundation for Life Sciences, Bioorganic Research Institute, Kyoto, Japan; Department of Chemistry, Graduate School of Science, The University of Osaka, Osaka, Japan

**Keywords:** iron, nicotianamine, small intestine, organoid, 2D monolayer

## Abstract

Iron absorption in the small intestine has classically been described by the duodenal DMT1/FPN1 pathway for inorganic non-heme iron, yet emerging evidence suggests that chemically distinct iron forms may use region-specific routes. Nicotianamine (NA), a plant-derived metal chelator, can form NA–iron (NA–Fe) complexes and has been proposed to support intestinal iron absorption through amino acid transporter pathways. However, direct comparisons of transepithelial transfer of inorganic iron and NA–Fe across defined small intestinal regions under controlled epithelial conditions remain limited. Here, we established region-specific 2D epithelial monolayers derived from duodenal and proximal jejunal crypt organoids from male ICR mice cultured on Transwell inserts. Transcriptomic profiling indicated partial retention of regional identity, and barrier integrity was confirmed by junctional marker localization, transepithelial electrical resistance, and low paracellular permeability. We then examined expression and polarized localization of candidate transporters for inorganic iron (Dmt1/Fpn1) and NA–Fe (Pat1/Lat2). Finally, we quantified transepithelial transport using apical loading of isotope-labeled iron (^55^Fe) or NA–^55^Fe and measured radioactivity appearing in the basolateral compartment as the primary readout of transepithelial flux. Basolateral appearance of inorganic ^55^Fe was comparable between duodenum- and proximal jejunum-derived monolayers, whereas NA–^55^Fe exhibited significantly greater basolateral appearance in proximal jejunum-derived monolayers. These findings demonstrate that organoid derived, region-specific monolayers provide a tractable epithelial platform to evaluate iron form–dependent, region-specific transepithelial transfer and to enable further mechanistic dissection of NA–Fe transport.

**NEW & NOTEWORTHY:** Non-heme iron absorption may depend on iron chemical form and intestinal region, but direct epithelial comparisons are scarce. We established duodenum and proximal jejunum derived murine intestinal organoid monolayers on Transwells and quantified transepithelial flux using isotope-labeled iron. Inorganic ^55^Fe showed no clear regional difference, whereas NA–^55^Fe displayed greater basolateral appearance in proximal jejunum-derived monolayers. This platform enables mechanistic studies of NA–iron complex transport.

## INTRODUCTION

Iron is an essential trace element that supports diverse physiological functions, including oxygen transport and DNA synthesis, and iron deficiency remains an important nutritional challenge(1). In mammals, systemic iron homeostasis is largely determined by intestinal absorption(2, 3). Classically, dietary non-heme iron is absorbed predominantly in the duodenum through a polarized epithelial transport process: luminal ferric iron is reduced to ferrous iron by the brush-border ferrireductase duodenal cytochrome b (Dcytb), taken up across the apical membrane via divalent metal transporter 1 (DMT1; SLC11A2), and subsequently exported across the basolateral membrane into the circulation via ferroportin 1 (FPN1; SLC40A1)(3–6). However, because dietary iron can exist in multiple chemical forms, iron absorption may not be fully explained by the duodenal DMT1/FPN1 axis alone(7).

Nicotianamine (NA) is an iron chelator ubiquitous in higher plants and is indispensable for iron transport in plants. Membrane transport of NA–Fe(II) complexes is mediated by yellow stripe 1/yellow stripe-like (YS1/YSL) transport proteins (8, 9). NA–Fe(II) plays an essential role in plant iron homeostasis, and plant foods such as legumes, wheat, and vegetables contain significant amounts of NA (10). Moreover, NA has been shown to strongly enhance iron uptake in the human colon carcinoma-derived Caco-2 monolayer model (11). Despite these observations, whether NA–Fe(II) directly contributes to iron absorption in animals and the molecular mechanisms by which NA–Fe(II) is taken up in the intestine has not been fully understood.

More recently, our group reported that NA–Fe(II) complex can be absorbed from the intestine as chelated complexes (12, 13). While inorganic iron is typically absorbed and exported in the duodenum, these findings supported a model in which NA–Fe(II) is taken up from the luminal side of proximal jejunal epithelium via the amino acid transporter PAT1 (SLC36A1) and exported across the basolateral membrane via the LAT2 (SLC7A8)/4F2hc (SLC3A2) heterodimer (hereafter referred to as LAT2) (12, 13). Collectively, this framework suggests that non-heme iron absorption may involve multiple pathways that depend on both iron chemical form and small intestinal region.

The small intestine exhibits regional specialization in both morphology and absorptive physiology. Along the proximal–distal axis, differences in epithelial organization, luminal conditions, and transporter expression contribute to functional differences between the duodenum, jejunum, and ileum (14, 15). Therefore, the intestinal handling of chemically distinct forms of non-heme iron may depend not only on transporter usage, but also on the region of the small intestine where transport occurs.

To elucidate iron form- and region-dependent transport mechanisms in this context, it is valuable to use physiologically relevant models that preserve epithelial polarity and barrier integrity while enabling quantification of transepithelial transfer. However, within the body, it is difficult to isolate epithelial-intrinsic transport processes because intestinal absorption is influenced by multiple factors, including luminal chemistry and the microbiota (14). Conversely, *in vitro* cell line models are experimentally tractable but may not adequately maintain regional identity or the expression and polarized localization of transporters. For example, Caco-2 monolayers have been widely used for trace element transport studies, yet concerns have been raised regarding their ability to reflect native small intestinal physiology and regional specificity (11, 16, 17).

Although organoid- or iPSC-derived intestinal epithelial cells have recently been adapted into 2D monolayers for transport assays, comparative analyses of transepithelial transfer across different small intestinal region remain limited (18–20). In this study, to evaluate non-heme iron transepithelial transport as a function of iron chemical form and small intestinal region, we generated region-specific 2D monolayers from murine duodenal and proximal jejunal organoids. Using apical administration of isotope-labeled iron and quantification of its appearance on the basolateral side as the primary readout of transepithelial flux, we sought to establish a platform that enables direct comparisons of iron transport between iron forms and small intestinal regions.

## MATERIALS AND METHODS

### Animals

Male ICR mice were purchased from CLEA Japan, acclimatized, and maintained on a standard feed (CE-2, CLEA Japan) with free access to water. All animal experiments were approved by the Suntory animal ethics committee (APRV000025) and were performed according to the institutional guidelines. Mice were euthanized by CO_2_ asphyxiation.

### Crypt isolation and 3D organoid formation

Crypt isolation and 3D organoid formation were performed as previously described (21–23). Briefly, proximal segments of the small intestine, including the duodenum (~5 cm distal to the pylorus) and the proximal jejunum (the upper half of the small intestine excluding the duodenum), were isolated separately (13). Subsequently, fat tissues attached to the small intestine were removed, and the lumen was flushed to remove luminal contents. The tissues were opened longitudinally and washed with cold phosphate-buffered saline (PBS) supplemented with 100 U/mL penicillin/streptomycin (Gibco) and 50 μg/mL gentamicin sulfate solution (Nacalai Tesque) (Antibiotic PBS; PBS-ABx). The tissues were cut into 5 × 5 mm pieces and further washed with cold PBS-ABx. Duodenal and proximal jejunal fragments were incubated separately in PBS-ABx containing 2 mM EDTA (Nacalai Tesque) for 30 min on ice. After removal of the EDTA solution, the fragments were vigorously shaken by hand in cold PBS-ABx to release crypt–villus complexes. The resulting suspensions were passed through a 70-μm cell strainer to remove residual villous material, and isolated crypts were centrifuged (390 × g for 3 min at 4 °C). The pellet was resuspended in Advanced DMEM/F-12 (Gibco) containing 2% (w/v) sorbitol (Nacalai Tesque) and 1% (v/v) FBS (Sigma-Aldrich), and low-speed centrifugation (80 × *g* for 3 min at 4 °C) was repeated twice to remove unwanted cells. Purified crypts (100 duodenal or proximal jejunal crypts) were mixed with 40 μL Matrigel (Corning) and plated per well in a pre-warmed 24-well plate (IWAKI). After Matrigel polymerization, 500 μL of 3D organoid culture medium (Advanced DMEM/F-12 supplemented with 50 U/mL penicillin/streptomycin, 25 μg/mL gentamicin, 2 mM L-glutamine (Gibco), 20 ng/mL mouse EGF(315-09, PeptoTech), 100 ng/mL mouse/rat Noggin (QK033, Qkine), and 150 ng/mL human R-spondin1 (QK006, Qkine)) was added. Organoids were maintained at 37 °C in a humidified atmosphere with 5% CO_2_, and the medium was changed every 2–3 days.

### Two-dimensional epithelial monolayer formation

Two-dimensional (2D) epithelial monolayer formation was performed as previously described (24). Prior to 2D monolayer formation, duodenal and proximal jejunal 3D organoids were maintained for 1–2 passages in 2D monolayer induction medium (IntestiCult Organoid Growth Medium for Human (ST-06010, STEMCELL Technologies) supplemented with 50 U/mL penicillin/streptomycin, 25 μg/mL gentamicin, 0.8 mM valproic acid (FUJIFILM Wako), 1% Wnt3a/Afamin CM (J2-005-05, MBL), and 10 μM Y-27632 (FUJIFILM Wako). For extracellular matrix (ECM) coating, 24-well translucent culture inserts (0.4 μm pore size, 1.0 × 10^8^ pores/cm^2^; 3413, Corning) were coated with 75 μL MatriMix-511 (nippi) according to the manufacturer’s instructions and incubated for 90 min at 37 °C. 3D organoids were collected from Matrigel and resuspended in cold PBS, followed by mechanical dissociation into single crypt domains. Samples were centrifuged (200 × g, 2 min, 4 °C), and the pellet was resuspended in 2 mL TrypLE Express (Gibco) and incubated at 37 °C for 5 min. Enzymatic dissociation was quenched by adding 2 mL Opti-MEM (Gibco), and cells were pelleted (300 × g, 2 min, 4 °C). The pellet was resuspended in 2 mL Opti-MEM and passed through a 30-μm cell strainer. Cell concentration was determined using a cell counter (CDA-1000, Sysmex, Japan). Samples were centrifuged once more (300 × g, 2 min, 4 °C) and resuspended in 2D monolayer induction medium (2.0–4.0 × 10^5^ cells in 100 μL per insert). After removing excess MatriMix solution from the inserts, 500 μL of 2D monolayer induction medium was added to the bottom chamber, and 100 μL of the cell suspension was gently applied to the upper insert. Monolayers were maintained at 37 °C in a humidified atmosphere with 5% CO_2_, and medium was changed every 2–3 days. For the experiments described below, monolayers cultured for 13–15 days were used, with 2D monolayer induction medium employed for the first 4–5 days, followed by 2D monolayer differentiation medium (a 1:1 mixture of 3D organoid culture medium and 2D monolayer induction medium, with Y-27632 adjusted to a final concentration of 10 μM).

### RNA sequencing

Total RNA was extracted from duodenal and proximal jejunal 2D monolayers and purified, including genomic DNA depletion, using the NucleoSpin RNA Plus XS kit (MACHEREY-NAGEL) according to the manufacturer’s instructions. Three independent biological replicates from each region were prepared for RNA sequencing (RNA-seq). Quality-controlled RNA (RIN > 8) was used to generate strand-specific, poly(A)-enriched libraries, which were sequenced as 150-bp paired-end reads by Novogene (Beijing, China) using an Illumina NovaSeq X Plus system (Illumina). The RNA-seq data generated in this study have been deposited in the NCBI Sequence Read Archive (SRA) under BioProject accession no. PRJNA1441194 (run accessions SRR37725115-SRR37725120) and will be made publicly available upon publication of this article. In addition to our dataset, public RNA-seq datasets for duodenum (GSE102789; SRA run accessions SRR5944370, SRR5944371, and SRR5944375) and proximal jejunum (GSE182348; SRA run accessions SRR15509804-SRR15509806) were included (25, 26). Reads were aligned to the mouse reference genome (mm10) using HISAT2 (v2.2.1), and transcript abundance was estimated as transcripts per million (TPM) using Cufflinks (v2.2.1). For visualization and inter-sample comparisons, TPM values were log2-transformed as log_2_(TPM + 1), where indicated.

### Immunofluorescence staining

Mouse small intestinal tissues (duodenum and proximal jejunum) and 2D monolayers were fixed in 4% paraformaldehyde (Nacalai Tesque) for 1–3 h at 4 °C, washed with PBS (pH 7.4, Gibco), and cryoprotected by sequential incubation in 15% and 30% (w/v) sucrose (Nacalai Tesque) in PBS. Samples were embedded in OCT compound (Sakura Finetek) and stored at −80 °C until cryosectioning. For 2D monolayers, monolayers were excised together with the membrane from the culture insert prior to embedding. Cryosections (8 μm) were prepared using a CryoStar NX70 cryostat (PHC epredia). Sections were air-dried for 30 min at 37 °C, rinsed with PBS (2 min), and permeabilized with PBS containing 0.2% Triton X-100 (Nacalai Tesque) (PBSTx) for 5 min. Sections were blocked with 1% bovine serum albumin (BSA; Sigma-Aldrich) in PBSTx for 60 min at room temperature. For triple immunofluorescence staining, sections were incubated overnight at 4 °C with a primary antibody mixture containing goat anti-Villin polyclonal antibody (1:100; sc-7672, Santa Cruz Biotechnology), Alexa Fluor 647–conjugated rat anti-EpCAM monoclonal antibody (1:200; 118211, BioLegend), and one of the following rabbit primary antibodies against iron-related transporters: anti-DMT1 (1:100; 20507-1-AP, Proteintech), anti-FPN1 (1:200; NBP1-21502, Novusbio), anti-PAT1 (1:50; NBP1-81280, Novusbio), or anti-LAT2 (1:100; ab75610, Abcam) (performed in separate sections). After washing, sections were incubated for 1 h at room temperature with DyLight 488-conjugated donkey anti-goat IgG antibody (1:1000; SA5-10086, Thermo Fisher) and Cyanine3-conjugated donkey anti-rabbit IgG antibody (1:1000; 40602, BioLegend) with Hoechst 33342 (2 µg/mL; Nacalai Tesque) for nuclear counterstaining. Following two additional PBS washes, sections were mounted using VECTASHIELD Vibrance Antifade Mounting Medium (Vector Laboratories) and covered with a glass coverslip (Matsunami). Imaging was performed using an FV3000 confocal laser scanning microscope (Olympus). Images were acquired sequentially to avoid spectral overlap.

### TEER Measurements

Transepithelial electrical resistance (TEER) of differentiated 2D monolayers was measured using a Millicell ERS-2 voltohmmeter (Merck Millipore) according to the manufacturer’s instructions. Measurements were performed at room temperature. TEER values (Ω·cm2) were calculated by subtracting the resistance of a blank insert (MatriMix-coated insert without cells) and multiplying by the membrane area (0.33 cm^2^).

### Lucifer yellow permeability assay

Culture medium was removed from both sides, and differentiated 2D monolayers were rinsed twice with the transport buffer (TB) consisting of Hank’s Balanced Salt Solution (HBSS, Nacalai Tesque) and 10 mM HEPES (pH 7.4, Gibco). The monolayers were preincubated for 15 min at 37 °C with TB added to the apical (100 μL) and basolateral (500 μL) chambers. After removal of the buffer, 100 μL of TB containing 60 μM Lucifer yellow CH dipotassium salt (FUJIFILM Wako) was added to the apical chamber, and 500 μL of TB was added to the basolateral chamber, respectively. After incubation for 90 min at 37 °C, samples were collected from the basolateral side and fluorescence was measured using a FlexStation 3 multi-mode microplate reader (Molecular Devices) at 428 nm excitation and 535 nm emission. Lucifer yellow concentrations were calculated from a standard curve generated by serial dilutions of Lucifer yellow. The apparent permeability coefficient (Papp) was calculated as:

Papp = (dCr/dt) × Vr / (A × C_0_),

where dCr/dt is the rate of change in receiver concentration, Vr is the receiver volume, A is the membrane area, and C_0_ is the initial donor concentration.

### NA synthesis and NA-Fe(II) complex preparation

The NA used in this study (purity > 95%, as determined by 1H NMR) was prepared by chemical synthesis based on previously reported methods for NA-related compounds (27, 28). The synthetic NA was subsequently used for preparation of the NA-Fe(II) complex for transport assays as previously described (12, 29, 30).

### Iron transport assay

Iron transport was assessed essentially as previously described (12). In brief, culture medium was removed from both chambers, and monolayers were washed twice with TB. The monolayers were preincubated for 15 min at 37 °C with TB added to the apical (200 μL) and basolateral (400 μL) chambers. After removal of the buffer, the apical chamber was loaded with 100 μL of TB containing 2 mM or 10 mM sodium L-ascorbic acid (Sigma-Aldrich) and either 1 mM ^55^Fe(II) (Fe-55 in 1M HCl 10 µg/ml, 0.903 Ci/g, radionuclidic Purity: > 99%; 6055 Eckert & Ziegler) adjusted to pH 7.4 with 1M NaOH or NA-^55^Fe(II) complex (10 mM NA:1 mM ^55^Fe, 10:1). The basolateral chamber was loaded with 400 μL of TB. After incubation for 30 min at 37 °C, apical buffer was aspirated and the basolateral buffer was collected into scintillation vials (986492, WHEATON) for counting. The monolayers were rinsed once with TB, excised from the insert, and transferred to a microcentrifuge tube containing glass beads and 200 µL of distilled water. Samples were vortexed for 1 min, centrifuged (5,000 × g, 3 min), and the supernatant was used as the cell lysate. Basolateral samples (300 μL) or cell lysates (200 μL) were mixed with 3 mL Ultima Gold scintillation cocktail (PerkinElmer) and radioactivity was measured using an ALOKA LSC-6100 liquid scintillation counter (Hitachi-Aloka Medical).

### Statistical analysis

Two-tailed unpaired Student’s t tests were used to compare data between two groups. Data and statistical analyses were performed using Microsoft Excel. Data are presented as the mean ± standard deviation (SD). In all cases, differences were considered significant when *p* < 0.05. All experiments were repeated at least three times.

## RESULTS

### Generation of region-specific murine small intestinal 2D monolayers and assessment of regional identity and cellular composition

Based on previous findings (12, 13) that the absorption of non-heme iron depends on the location in the small intestine (SI), which is influenced by the chemical form of iron (inorganic iron vs. NA–Fe complexes), crypts from the duodenum (Duod) and proximal jejunum (ProxJej) were isolated from male ICR mice maintained under standard housing conditions. Using isolated crypts, we established 3D organoids and subsequently generated region-specific 2D epithelial monolayers on Transwell inserts (Fig. 1A)(24). To evaluate the cellular composition and regional identity of the induced monolayers, RNA sequencing (RNA-seq) was performed, and the transcriptomes were compared with publicly available *in vivo* small intestinal datasets (Duod: GSE102789; ProxJej: GSE182348) (Fig. 1B–H)(25, 26). Analysis of the expression of small intestinal cell markers revealed that the expression level of *Lgr5*, a marker for intestinal stem cells, was higher in the Duod- and ProxJej-2D monolayers than Duod- and ProxJej-SI (Fig. 1B). Furthermore, the expression of the absorptive epithelial marker *Vil1* was comparable between SI and 2D conditions (Fig. 1C). However, the expression of secretory lineage markers (*Lyz1, Chga*, and *Muc2*) was lower in the 2D monolayers (Fig. 1D–F). These results suggest that the monolayers generated under our conditions are mainly composed of intestinal stem cells and absorptive epithelial cells (enterocytes).

**Figure 1.**
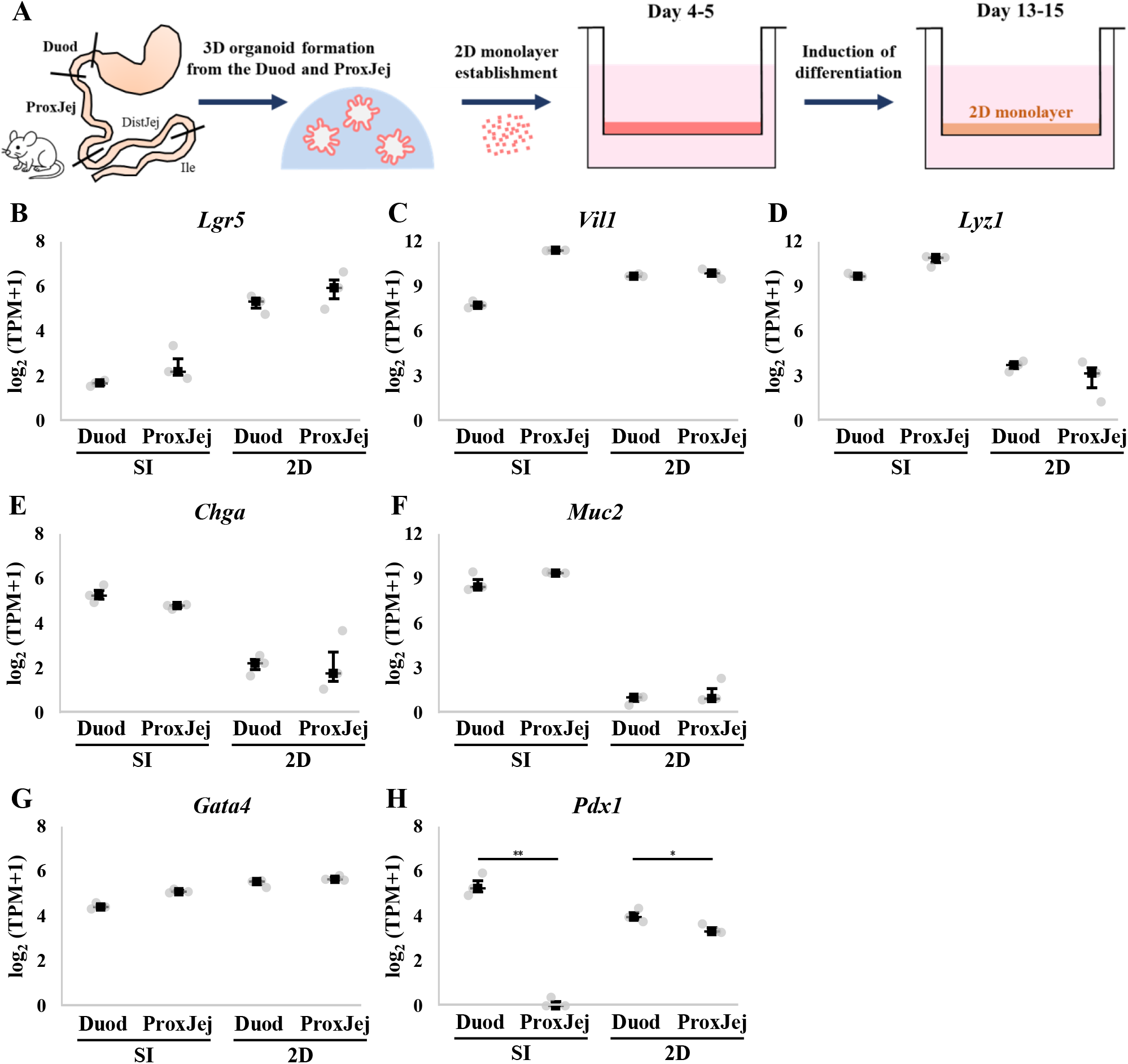
Generation of region-specific murine small intestinal 2D monolayers and assessment of regional identity and cellular composition. (A) Schematic overview of the experimental workflow. Crypts were isolated from the Duod and ProxJej of ICR mice to establish 3D organoids. Single cells derived from the 3D organoids were seeded onto Transwell inserts, and a 2D epithelial monolayer was formed through stepwise induction of differentiation. (B–F) RNA-seq expression of intestinal cell-type marker genes in region-specific 2D monolayers and SI (public datasets: Duod, GSE102789; ProxJej, GSE182348). n=3 independent biological samples. Points, samples; square, median; vertical bars, IQR (Q1–Q3), values are log_2_(TPM+1). (B) intestinal stem cell marker *Lgr5*; (C) absorptive epithelial marker *Vil1*; (D) Paneth cell marker *Lyz1*; (E) enteroendocrine cell marker *Chga*; (F) goblet cell marker *Muc2*. (G, H) RNA-seq expression of small intestinal regional marker genes: proximal small intestinal marker *Gata4* (G) and duodenal marker *Pdx1* (H). Duod, duodenum; ProxJej, proximal jejunum; DistJej, distal jejunum; Ile, ileum; SI, small intestine; IQR, interquartile range. \**p* < 0.05, \*\**p* < 0.01.

We next examined the expression of intestinal regional marker genes (31, 32). The expression of the proximal SI marker *Gata4* was comparable between Duod and ProxJej in both conditions (Fig. 1G). The expression of the duodenum marker *Pdx1* was higher in the Duod monolayer than in the ProxJej monolayer (Fig. 1H). Although the regional differences were less pronounced than those observed in SI, the region-specific 2D monolayers retained their regional characteristics.

### Barrier function of region-specific 2D monolayers

Immunostaining for ZO-1 and E-cadherin confirmed stable formation of tight junctions and adherens junctions in both Duod- and ProxJej-2D monolayers (Fig. 2A, B). TEER values (mean ± SD) were 152.2 ± 13.7 Ω·cm^2^ for Duod-2D and 136.2 ± 6.4 Ω·cm^2^ for ProxJej-2D (Fig. 2C, n = 6 each), which were comparable to or higher than the reported value for murine small intestine (33). In addition, the apparent permeability coefficient (Papp) of Lucifer Yellow (MW = 521) was low in both monolayers (0.07 ± 0.037 × 10^−6^ cm/s for Duod-2D and 0.19 ± 0.067 × 10^−6^ cm/s for ProxJej-2D; Fig. 2D, n = 6 each). Collectively, these data indicate that both region-specific monolayers possess sufficient barrier function for transepithelial transport assays and allow comparisons between intestinal regions.

**Figure 2.**
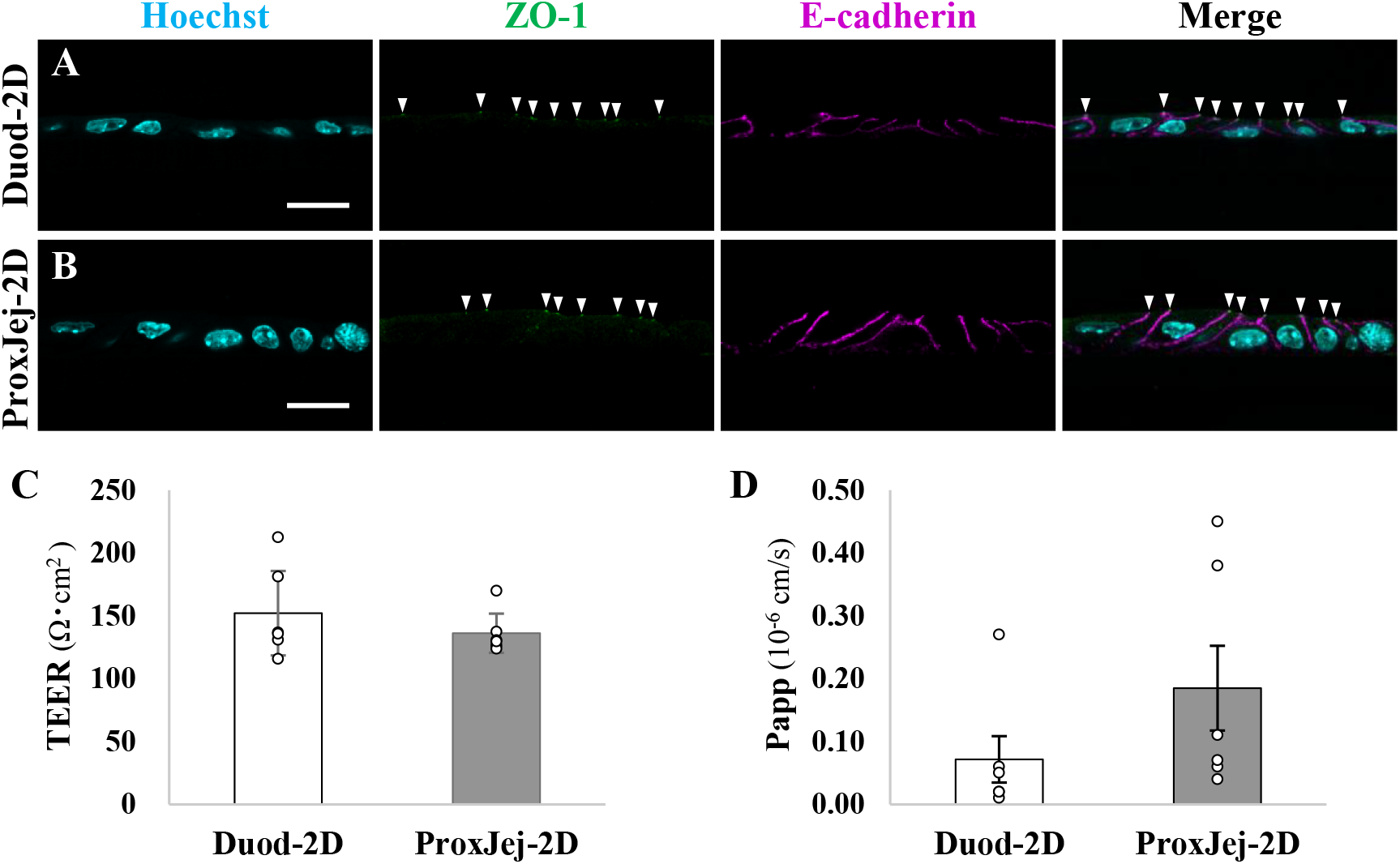
Barrier function of region-specific 2D monolayers. (A, B) Immunofluorescence images of cryosections of Duod-2D (A) and ProxJej-2D monolayers (B). Nuclei were stained with Hoechst; TJs were labeled with ZO-1 (arrowheads); AJs were labeled with E-cadherin. (C) TEER of each 2D monolayer (n = 6 each). (D) Papp for LY in each 2D monolayer (n = 6 each). The graphs present the mean ± SD. Scale bars, 20 µm. Duod, duodenum; ProxJej, proximal jejunum; TJ, tight junction; AJ, adherens junction; TEER, transepithelial electrical resistance; Papp, Apparent permeability coefficient; LY, lucifer Yellow.

### Expression and localization of iron/NA–iron-related transporters in region-specific 2D monolayers

Before performing transepithelial transport assays for Fe(II) and NA–Fe(II) complex, we examined mRNA expression (RNA-seq) and protein localization (IHC) of their related transporters (Fig. 3). The apical transporter *Dmt1 (Slc11a2)* for non-heme iron was expressed at similar levels in Duod- and ProxJej-SI, as well as in these region-derived 2D monolayers (Fig. 3A, B). In contrast, the basolateral export transporter *Fpn1 (Slc40a1)* was highly expressed in Duod-SI compared to ProxJej-SI, but was comparably expressed in 2D monolayers (Fig. 3A, C). It has been reported that under normal feeding (iron-sufficient) conditions, Dmt1 shows weak apical localization in the intestinal epithelium (34). Consistent with this, IHC for Dmt1 in SI revealed little detectable signal at the apical side of epithelial cells in both regions (Fig. S1A, B). Although the Fpn1 signal in SI did not show a clear regional difference comparable to the *mRNA* expression (Fig 3C), Fpn1 protein was observed along the basolateral membrane (Fig. S1C, D). In the 2D monolayers, Dmt1 protein was detected at the apical surface in both regions, but the signal was relatively weaker in ProxJej-2D (Fig. 3D, E). Fpn1 localization was observed at the basolateral membrane in both regions (Fig. 3F, G).

**Figure 3.**
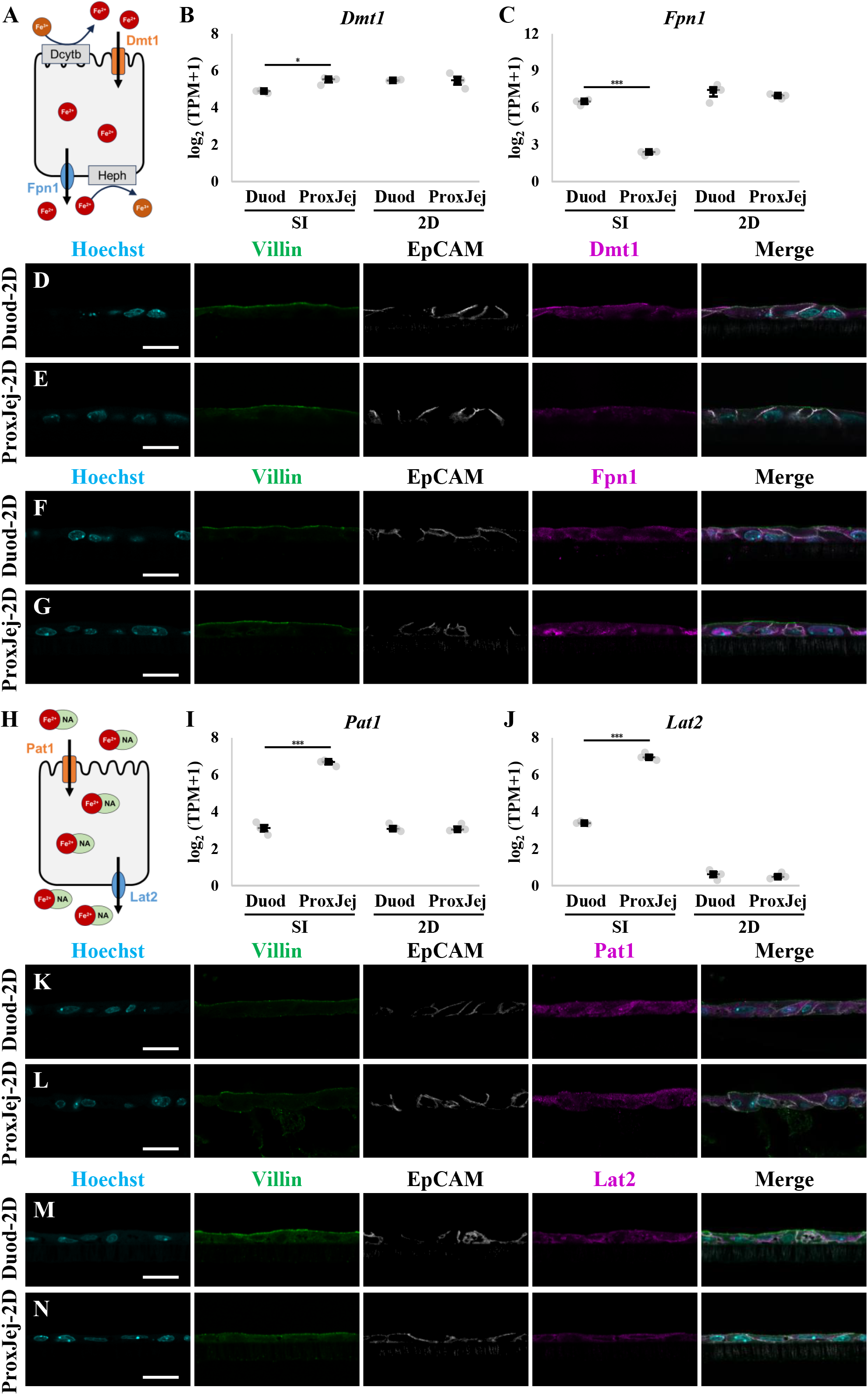
Expression and localization of iron/NA–iron-related transporters in region-specific 2D monolayers. (A) Schematic model of inorganic iron uptake and export in the enterocytes. Ferrous iron (Fe(II)), which is reduced from Fe(III) by Dcytb, is taken up via Dmt1 and exported via Fpn1. The exported Fe(II) is oxidized to Fe(III) by Heph. (B, C) RNA-seq expression of *Dmt1* (B) and *Fpn1* (C) in region-specific 2D monolayers and SI. n=3 independent biological samples. Points, samples; square, median; vertical bars, IQR (Q1–Q3), values are log_2_(TPM+1). (D-G) Immunofluorescence images of Dmt1 (D, E) and Fpn1 (F, G) in cryosections of Duod- and ProxJej-2D monolayers. Nuclei were stained with Hoechst; Villin and EpCAM were used as apical and basolateral markers, respectively. (H) Schematic model of NA–iron complex uptake and export in the enterocytes. NA-Fe(II) is taken up via Pat1 and exported via Lat2. (I, J) RNA-seq expression of *Pat1* (I) and *Lat2* (J) in region-specific 2D monolayers and SI. (K-N) Immunofluorescence images of Pat1 (K, L) and Lat2 (M, N) in cryosections of Duod- and ProxJej-2D monolayers. Dcytb, duodenal cytochrome b; Heph, hephaestin; Duod, duodenum; ProxJej, proximal jejunum; SI, small intestine; IQR, Interquartile Range, NA, nicotianamine. \**p* < 0.05, \*\*\**p* < 0.005. Scale bars, 20 µm.

For NA–Fe(II) transport, *Pat1 (Slc36a1)* and *Lat2 (Slc7a8)* showed higher mRNA expression in ProxJej-SI than in Duo-SI, whereas region-dependent differences were less apparent in the 2D monolayers (Fig. 3H-J). Protein localization analysis further showed apical Pat1 and basolateral Lat2 signals in both SI and 2D monolayers (Figs. 3K–N and S2). However, regional differences in IHC signal were less pronounced than those observed in *mRNA* expression.

### Transepithelial flux of isotope-labeled iron across region-specific 2D monolayers

To evaluate transepithelial flux of inorganic iron and NA–iron complexes across the 2D monolayers, we loaded transfer buffer (TB) containing 2 mM ascorbic acid and 1 mM inorganic ^55^Fe(II) or NA–^55^Fe(II) complex to the apical chamber, and TB alone to the basolateral chamber. After 30 min, radioactivity in the basolateral solution was measured (Fig. 4A). With inorganic ^55^Fe(II), basolateral radioactivity (mean ± SD) was similar between Duod-2D (20.85 ± 1.91 counts/5 min) and ProxJej-2D (23.47 ± 2.81 counts/5 min) (Fig. 4B, n = 5 each). In contrast, with NA–^55^Fe(II) complex, basolateral radioactivity was significantly higher in ProxJej-2D (52.80 ± 18.51 counts/5 min) than in Duod-2D (20.05 ± 9.19 counts/5 min) (Fig. 4B, n = 5 each). We also quantified intracellular radioactivity and found that, with inorganic ^55^Fe(II), no regional differences were observed under 2 mM ascorbic acid conditions; however, under 10 mM ascorbic acid conditions, Duod-2D tended to exhibit higher intracellular radioactivity than ProxJej-2D (Fig. S3). Together, these results suggest that the region-specific 2D monolayer platform recapitulates, at least in part, small intestinal region dependent non-heme iron transport, which depends on iron chemical form (inorganic iron vs NA–iron complex), similar to the *in vivo* situation (12, 13).

**Figure 4.**
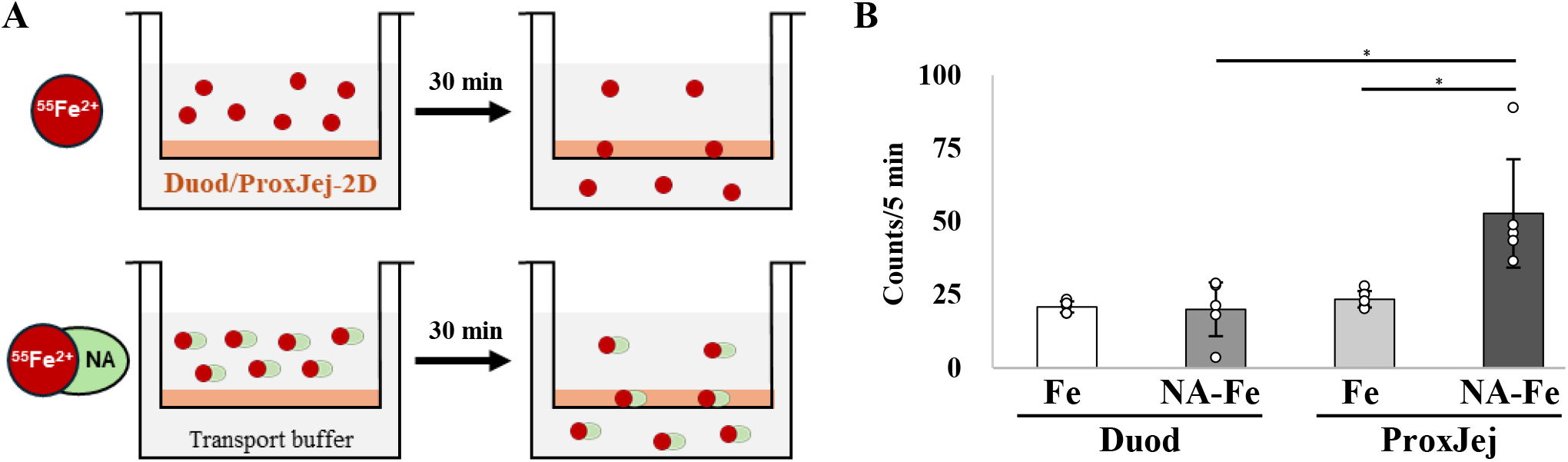
Transepithelial flux of isotope-labeled iron across region-specific 2D monolayers. (A) Schematic overview of the iron transport assay. Isotope-labeled inorganic iron (^55^Fe(II)) or NA–^55^Fe(II) complex was added to the apical chamber of each region-specific 2D monolayer cultured on Transwell inserts, and radioactivity appearing in the basolateral chamber was quantified after 30 min incubation. (B) Basolateral appearance of ^55^Fe per 5 min in 2 mM As and 1 mM ^55^Fe or 1 mM NA-^55^Fe (NA:Fe=10:1) conditions (n = 5 each). The graphs present the mean ± SD. Duod, duodenum; ProxJej, proximal jejunum; NA, nicotianamine; As, ascorbic acid. \**p* < 0.05.

## DISCUSSION

In this study, we aimed to establish a region-specific murine small intestinal 2D monolayer platform that enables direct comparisons of transepithelial transport properties, which depend on the non-heme iron chemical form (Fe(II) vs NA–Fe(II) complexes) under controlled conditions without confounding influences from luminal contents or the microbiota. Using 2D monolayers derived from duodenal (Duod) and proximal jejunal (ProxJej) organoids from ICR mice (Fig. 1), isotope tracer assays showed that basolateral transfer of ^55^Fe(II) was not markedly different between regions, whereas NA–^55^Fe(II) exhibited significantly greater basolateral appearance (transepithelial flux) in ProxJej-2D (Fig. 4). These findings support the emerging framework that non-heme iron absorption is not mediated by a single pathway, but can involve distinct intestinal regions and molecular mechanisms depending on the iron chemical form. In addition, our platform provides an experimental basis for further mechanistic studies aimed at determining transporter dependence, rate-limiting steps, and regulatory mechanisms for NA–Fe(II) transport.

An important observation in this study is that iron/NA-iron-related transporter mRNA expression patterns obtained by RNA-seq did not fully match those reported for the small intestine (SI) (Fig. 3) (12, 13), whereas the functional readout of transepithelial flux was consistent with the trends suggested by *in vivo* studies (Fig. 4) (13). This dissociation between the phenotype (transport function) and mRNA expression likely reflects the differences in differentiation state and cell-type composition of 2D monolayers. Regional marker genes (*Pdx1* and *Gata4*) indicated partial retention of the original tissue identity (Fig. 1G, H); however, the monolayers were enriched in stem-like and absorptive populations, showing limited secretory differentiation (Fig. 1B-F). This may dampen regional differences in transporter expression. More broadly, regional differences in the small intestine are shaped not only by transporter expression itself but also by tissue architecture and physiological context. *In vivo*, region-specific features such as epithelial maturation state, villus organization, luminal pH, and the microbiota are all likely to influence nutrient handling (14). Our 2D monolayer system does not fully reproduce all of these features, particularly the three-dimensional structure and the full diversity of differentiated epithelial cell types (Fig. 1). Nevertheless, the observation that NA–Fe(II) flux remained higher in ProxJej-2D (Fig. 4) suggests that at least part of the region-specific transport program is epithelial intrinsic and retained after organoid derivation. Additionally, further optimization of differentiation conditions may bring mRNA expression patterns closer to those observed *in vivo*.

Protein localization by IHC was generally consistent with prior knowledge in that uptake transporters localized to the apical membrane and export transporters localized to the basolateral membrane, with Dmt1 representing an exception (Fig. 3). Under normal feeding (iron-sufficient) conditions, intestinal Dmt1 apical localization is reported to be weak (34). Similarly, we observed little detectable apical Dmt1 signal in SI cryosections (Fig. S1A, B). In contrast, Dmt1 was clearly detected at the apical surface in the 2D monolayers, particularly in Duod-2D (Fig. 3D, E). This difference suggests that the monolayers may not be in an “iron-sufficient” state comparable to the *in vivo* duodenal epithelium. Further work assessing and manipulating cellular iron status (e.g., iron-responsive gene expression and ferritin levels) may help tune the monolayers toward a more physiological state.

In the iron transport assays, we used radioactivity appearing in the basolateral compartment after apical loading as the primary readout of transepithelial flux (Fig. 4). Using this readout, NA-Fe(II) exhibited a clear preference for ProxJej-2D (Fig. 4B). In contrast, regional differences were not evident in basolateral Fe(II) transfer, although intracellular radioactivity (a secondary readout reflecting uptake and retention) tended to be higher in Duod-2D under high ascorbate (10 mM) conditions (Figs. 4B, S3). One possible explanation is that increased ascorbate stabilizes iron in its ferrous state. This occurs by supporting reductive processes, such as Dcytb, and through ascorbate’s own reducing capacity (3, 5). Consequently, the availability of Fe^2+^ for Dmt1-mediated uptake increases. Moreover, enhanced apical uptake does not necessarily lead to a proportional increase in basolateral transfer. Previous study using Caco-2 monolayers has suggested that inorganic iron can be taken up intracellularly, whereas basolateral export is relatively low compared to chelated iron (12). This indicates that the efficiency of basolateral transfer may be influenced by chelation status.

Despite the lack of clear regional differences in *Pat1* and *Lat2* mRNA expression or IHC signal intensity in the 2D monolayers (Fig. 3I-N), NA–Fe(II) flux remained markedly higher in ProxJej-2D (Fig. 4B). Although the underlying factor(s) cannot be determined from the current data, it is possible that the rate-limiting step is not Lat2 expression itself, but rather heterodimer formation with 4F2hc, membrane localization, and/or transporter activity (13). In line with this possibility, *4F2hc* transcript abundance was similar between Duod-2D and ProxJej-2D in our RNA-seq dataset. However, whether protein abundance or heterodimer assembly differs between regions remains to be determined. Additionally, parallel pathways have been identified for other nutrients and metabolites beyond the initially presumed primary route, often in a region- or condition-dependent manner (35). Therefore, it is possible that, in addition to the Pat1-Lat2 pathway, other, as yet unknown, parallel mechanisms are involved in NA-Fe(II) transport. Functional analyses using Pat1- or Lat2-knockdown/knockout monolayers will be useful for directly testing these possibilities.

Finally, our region-specific, organoid-derived monolayer approach provides an alternative to conventional cell line models for evaluating region-dependent transepithelial transport. This approach offers a common experimental framework and enables genetic perturbations through engineered mouse knockout lines or organoid-based genome editing. Key remaining challenges include improving epithelial maturation, controlling cellular iron status for a better understanding of the uptake/export balance, and establishing transporter dependence through genetic or pharmacological perturbation.

## DATA AVAILABILITY

The data supporting the findings of this study are available in the article and its supplemental materials. The RNA-seq data generated in this study have been deposited in the NCBI Sequence Read Archive (SRA) under BioProject accession no. PRJNA1441194 (run accessions SRR37725115-SRR37725120) and will be released upon publication of this article.

## SUPPLEMENTAL MATERIAL

Supplemental Figs. S1–S3: https://doi.org/10.5281/zenodo.19228751

## ACKNOWLEDGMENTS

We thank Drs. Hiroyuki Yasui, Hiroyuki Kimura (currently at Kyoto University) and Hidekazu Kawashima from Kyoto Pharmaceutical University for their discussions and for the use of radioisotopes at the Radioisotope Research Center. We are grateful to Dr. Fumihiko Sato for valuable discussions.

## AUTHOR CONTRIBUTIONS

Y.T. designed and performed the experiments. Y.T., Y.M. and T.T wrote the original manuscript. Y.M. conducted the iron transport assay. K.N. carried out NA synthesis. All authors revised the manuscript.

## Supplemental Information

**Supplemental Figure S1.**
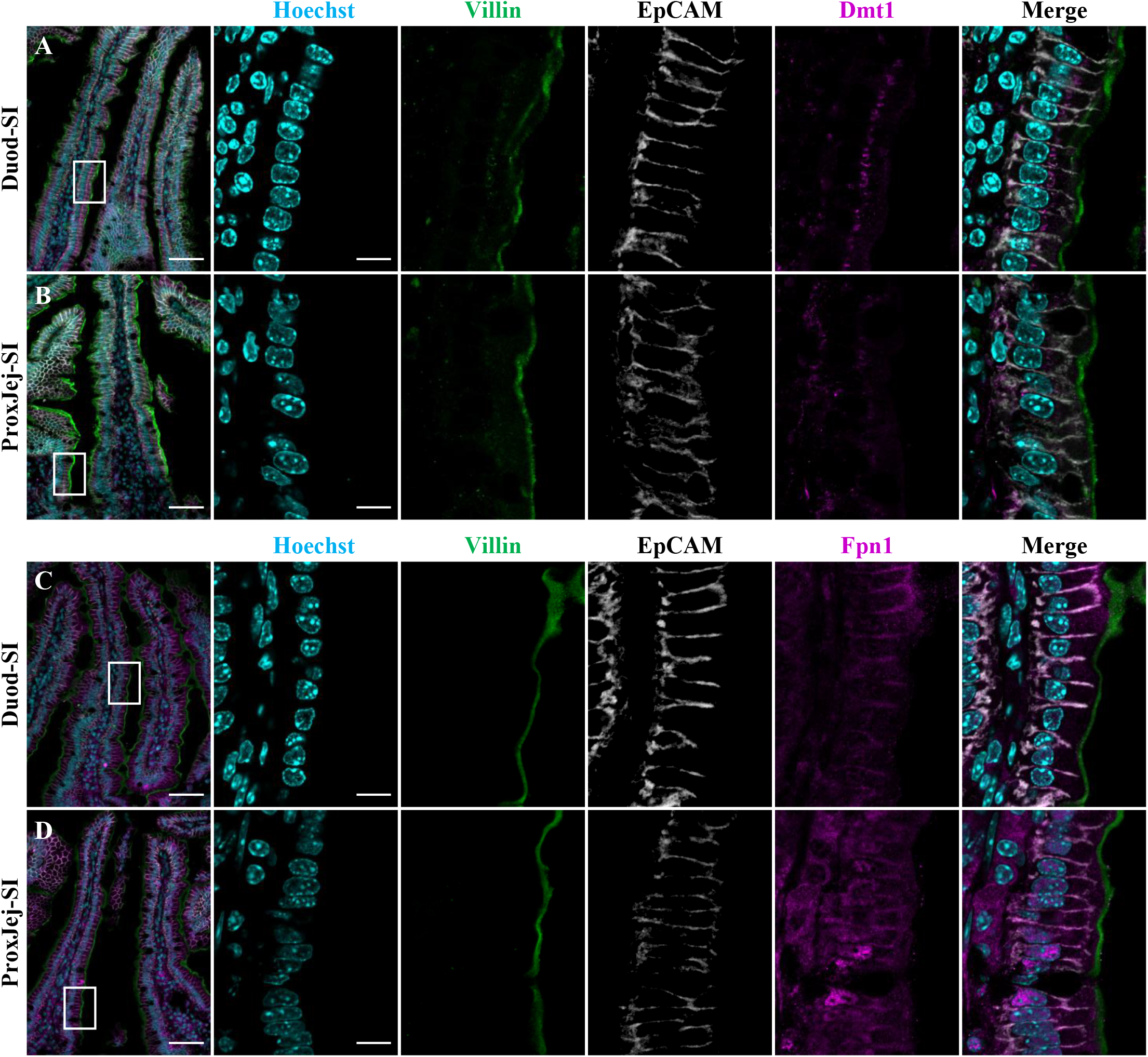
Localization of iron transporters in the murine duodenum and proximal jejunum. (A, B) Immunofluorescence images of Dmt1 in cryosections of Duod-SI (A) and ProxJej-SI (B). (C, D) Immunofluorescence images of Fpn1 in cryosections of Duod-SI (C) and ProxJej-SI (D). The right four images are enlarged views of the white squares in the corresponding left images. Nuclei were stained with Hoechst; Villin and EpCAM were used as apical and basolateral markers, respectively. Duod, duodenum; ProxJej, proximal jejunum; SI, small intestine. Scale bars, 50 μm (left); 10 μm (enlarged view).

**Supplemental Figure S2.**
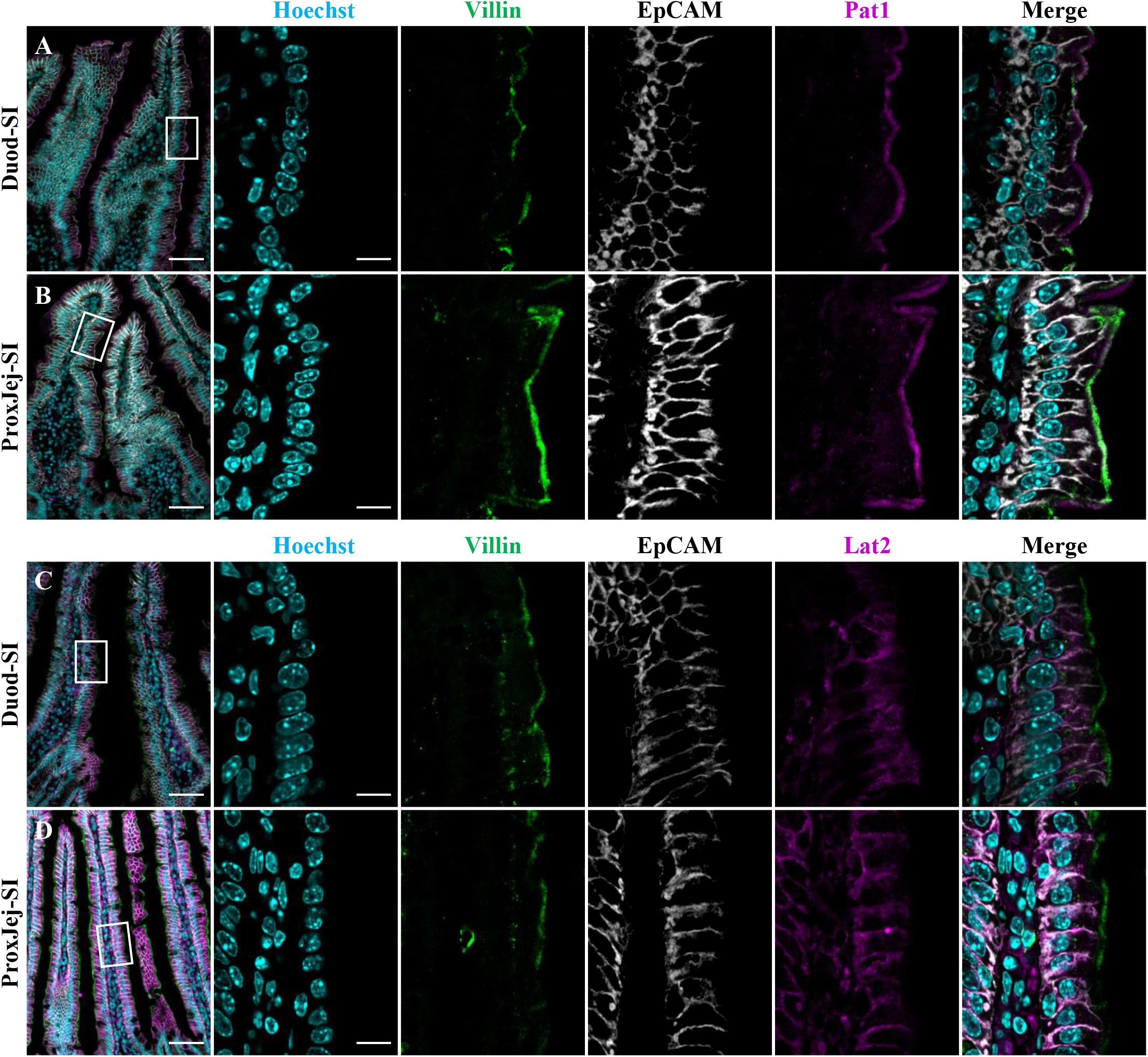
Localization of NA-iron transporters in the murine duodenum and proximal jejunum. (A, B) Immunofluorescence images of Pat1 in cryosections of Duod-SI (A) and ProxJej-SI (B). (C, D) Immunofluorescence images of Lat2 in cryosections of Duod-SI (C) and ProxJej-SI (D). The right four images are enlarged views of the white squares in the corresponding left images. Nuclei were stained with Hoechst; Villin and EpCAM were used as apical and basolateral markers, respectively. Duod, duodenum; ProxJej, proximal jejunum; SI, small intestine. Scale bars, 50 μm (left); 10 μm (enlarged view).

**Supplemental Figure S3.**
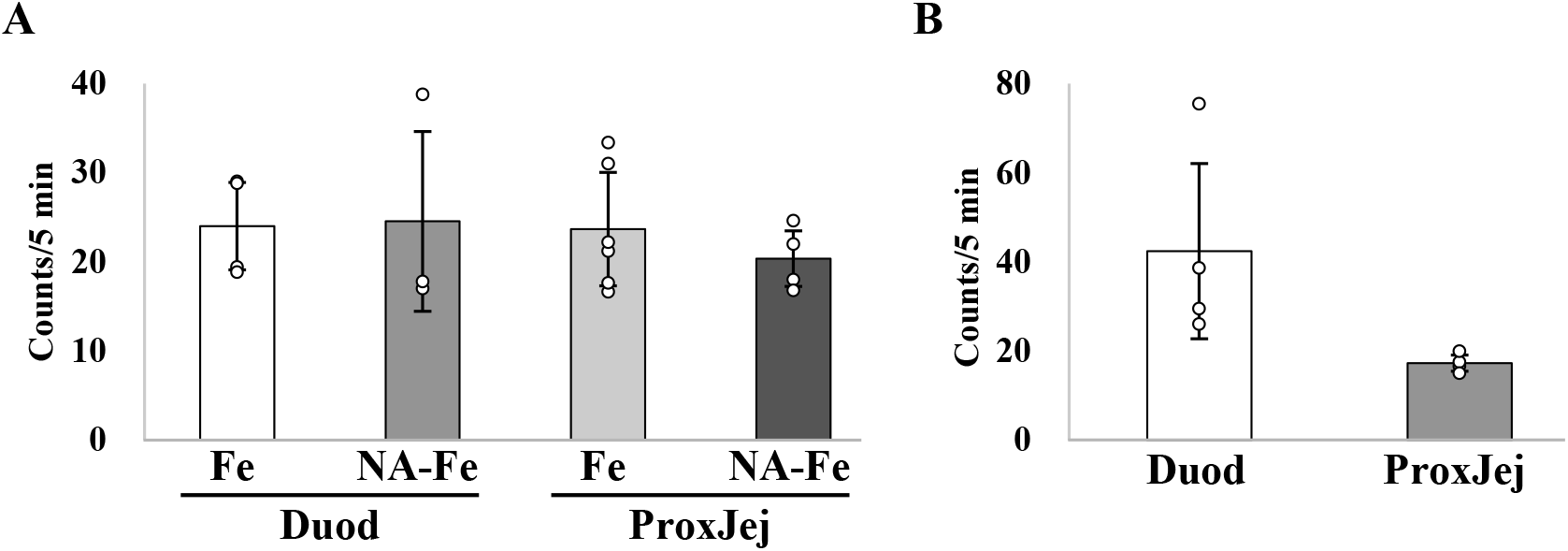
Intracellular accumulation of isotope-labeled iron across region-specific 2D monolayers. (A) Intracellular accumulation of ^55^Fe per 5 min in 2 mM As and 1 mM ^55^Fe or 1 mM NA-^55^Fe (NA:Fe=10:1) conditions (n = 3-6). (B) Intracellular accumulation of ^55^Fe per 5 min in 10 mM As and 1 mM ^55^Fe conditions (n = 3 each). The graphs present the mean ± SD. Duod, duodenum; ProxJej, proximal jejunum; NA, nicotianamine; As, ascorbic acid.

